# Integrated single-cell and spatial analysis identifies context-dependent myeloid-T cell interactions in head and neck cancer immune checkpoint blockade response

**DOI:** 10.1101/2025.03.24.644582

**Authors:** Athena E. Golfinos-Owens, Taja Lozar, Parth Khatri, Rong Hu, Paul M. Harari, Paul F. Lambert, Megan B. Fitzpatrick, Huy Q. Dinh

**Affiliations:** McArdle Laboratory for Cancer Research, University of Wisconsin School of Medicine and Public Health, Madison, WI 53792; University of Ljubljana, Ljubljana, Slovenia; Department of Surgical Oncology, Institute of Oncology Ljubljana, Ljubljana Slovenia; Department of Biostatistics and Medical Informatics, University of Wisconsin School of Medicine and Public Health, Madison, WI 53792; Department of Pathology and Laboratory Medicine, University of Wisconsin School of Medicine and Public Health, Madison, WI 53792; Department of Human Oncology, University of Wisconsin School of Medicine and Public Health, Madison, WI 53792

**Keywords:** scRNA-seq, spatial transcriptomics, head and neck squamous cell carcinoma, cellular neighborhoods, cell-cell interactions, ligand-receptor interactions, myeloid cells, immune checkpoint blockade

## Abstract

**Background:** Approximately 15-20% of head and neck cancer squamous cell carcinoma (HNSCC) patients respond favorably to immune checkpoint blockade (ICB). Previous single-cell RNA-Seq (scRNA-Seq) studies identified immune features, including macrophage subset ratios and T-cell subtypes, in HNSCC ICB response. However, the spatial features of HNSCC-infiltrated immune cells in response to ICB treatment need to be better characterized.

**Methods:** Here, we perform a systematic evaluation of cell interactions between immune cell types within the tumor microenvironment using spatial omics data using complementary techniques from both 10X Visium spot-based spatial transcriptomics and Nanostring CosMx single-cell spatial omics with RNA gene panel including 435 ligands and receptors. In this study, we used integrated bioinformatics analyses to identify cellular neighborhoods of co-localizing cell types in single-cell spatial transcriptomics and proteomics data. In addition, we used both publicly available scRNA-Seq and in-house spatial RNA-Seq data to identify spatially constrained Ligand-Receptor interactions in Responder patients.

**Results:** With 522,399 single cells profiled with both RNA and protein from 26 patients, in addition to spot-resolved spatial RNA-Seq from 8 patients treated with ICB together with bioinformatics analysis of publicly available single-cell and bulk RNA-Seq, we have identified a spatial and cell-type specific context-dependency of myeloid and T cell interaction difference between Responders and Non-Responders. We defined further cellular neighborhood and the sources of chemokine CXCL9/10-CXCR3 interactions in Responders, emerging targets in ICB, as well as CXCL16-CXCR6, CCL4/5-CCR5, and other underappreciated and potential markers and targets for ICB response in HNSCC. In addition, we have contributed a rich data resource of cell-cell Ligand Receptor interactions for the immunotherapy and HNSCC research community.

**Discussion:** Our work provides a comprehensive single-cell and spatial atlas of immune cell interactions that correlate with response to ICB in HNSCC. We showcase how integrating multiple technologies and bioinformatics approaches can provide new insights into potential immune-based biomarkers of ICB response. Our results suggested refining future studies using preclinical animal models in a more context-specific manner to elucidate potential underlying mechanisms that lead to improved ICB responses.

**What is already known on this topic:** Most cancer patients still do not experience clinical benefits from immune checkpoint blockade (ICB), necessitating the development of response biomarkers and new immunotherapeutic targets.

**What this study adds:** Here, we use integrated high-dimensional omics and bioinformatics approaches to identify immune cell-cell interaction markers associated with ICB response in patients with Head and neck squamous cell carcinoma.

**How this study might affect research, practice or policy:** We identified spatial and cell-type specificity of Ligand-Receptor interactions between myeloid and T cells in ICB Responder patients that may help inform further mechanistic studies and biomarker development.

## Introduction

Metastatic and recurrent head and neck squamous cell carcinoma (HNSCC) shows approximately 15-20% response rate to immune checkpoint blockade (ICB) therapy^1,2^. Understanding of the tumor microenvironment (TME) heterogeneity in HNSCC has rapidly increased due to the advance of high-dimensional single-cell omics data, allowing for the characterization of cellular compartments such as tumor and stromal cells^3,4^, immune cells^5^, and B cells^6^. Recent studies investigating ICB response in HNSCC have described valuable TME features, including the prognostic association of macrophage^7^, T cell^8^, and immune cell densities and tumor proximity of tertiary lymphoid structures markers^9^. Spatial omics and imaging technologies have identified HNSCC spatial TME features associated with survival^10^, human papillomavirus infection status^11^, invasive tumor phenotype^12^, and CD8+ T cell infiltration and dysfunction^13^. These insights have provided potential markers of ICB response. However, spatial and single-cell analysis of immune cell-cell interactions, which could potentially identify immunotherapeutic markers and targets for ICB treatment, is still lacking.

A scRNA-Seq analysis of HNSCC TME identified CXCL9:SPP1 macrophage polarization as a new marker for pro- and anti-tumoral tumor-associated macrophages, and interactions between myeloid and T cells, such as CXCL9-CXCR3 and CXCL16-CXCR6 associated with anti-tumoral features^7^. To systematically evaluate immune cell-cell interactions, we have generated spatial multi-omics data using 10X Visium and Nanostring CosMX platforms, together with a reanalysis of publicly available scRNA-seq^8^, to identify spatially-dependent immune features, such as cellular neighborhoods of co-localizing immune cell types, and Ligand-Receptor interaction pairs in HNSCC tumors from ICB-treated patients. We identified CD45+ cellular niches associated with ICB response, including higher frequencies of stromal-immune interface enriched for antigen-presenting and inflammatory cancer associated fibroblast (CAFs), dendritic cells (DCs) and CD8+ T cells in Responders to ICB. Furthermore, we defined a spatially aware and cell-type-specific context-dependent Ligand-Receptor pairs in different groups of ICB responses. This work provides a comprehensive single-cell and spatial atlas of immune cell interactions correlating with response to ICB in HNSCC and showcases how integrating multiple technologies and bioinformatics approaches can offer new insights into potential immune-based biomarkers of ICB response.

## Materials and Methods

A full description of the method is supplied in the Online Supplementary Methods section.

### In Brief

Nanostring CosMX single-cell spatial omics (protein and RNA) profiling was used for three HNSCC TMAs consisting of 68 cores from 24 patients (11 Responders and 13 Non-Responders), followed by integrated bioinformatics analysis with 8 10X Visium samples (4 Responders vs. 4 Non-Responders) using Seurat framework. Cellular neighborhood identification was based on a window-based approach^14^. In addition, we reanalyzed scRNA-Seq from treatment-naïve^5^ and ICB treatment^8^ with unified annotations of myeloid and T-cell subsets that allowed for cell-cell Ligand-Receptor interaction analysis using CellChat^15^, which was also utilized for CosMX RNA data. Other computational and statistical analysis were done in Python and R, details in Supplementary Methods.

## Results

### Multi-omics single-cell and spatial analyses of cellular neighborhood and Ligand-Receptor immune cell-cell interactions of HNSCC tumor microenvironment (TME) in response to ICB

To evaluate the spatially resolved immune landscape of ICB-treated HNSCCs, we collected Nanostring CosMX single-cell spatial transcriptomics and proteomics data from 68 tumor microarray (TMA) cores from 24 ICB-treated HNSCC patients (11 Responders and 13 Non-Responders) (**Fig 1A-B**). With 64 immune-centric protein markers, we identified most of the primary immune cell types, including three T cell subsets (CD4+, CD8+, and regulatory (Tregs) T cells), four myeloid cell types (monocytes, macrophages, dendritic cells - DCs, and neutrophils), together with B, plasma and NK cells with distinct lineage markers (**Fig 1A**) with 39.9% of cells, on average, are immune cells for both Responder and Non-Responder samples (**Fig 1A**). Except for tumor cells (Beta-catenin+PanRAS+EPCAM+), which decreased in Responders (p<0.01, **Fig 1A**), we observed trending changes in cell type frequencies between Responder and Non-Responder (CD8+ T cells higher in Responders, **Suppl Fig 1A**). With a 1000-gene RNA panel, we identified a larger extent of cell type identity, including three cancer-associated fibroblasts (CAF) subsets (myofibroblast, inflammatory, and antigen-presenting CAFs), and different tumor cell types expressing different keratin genes (**Fig 1B, Suppl Fig 1B**).

**Figure 1.**
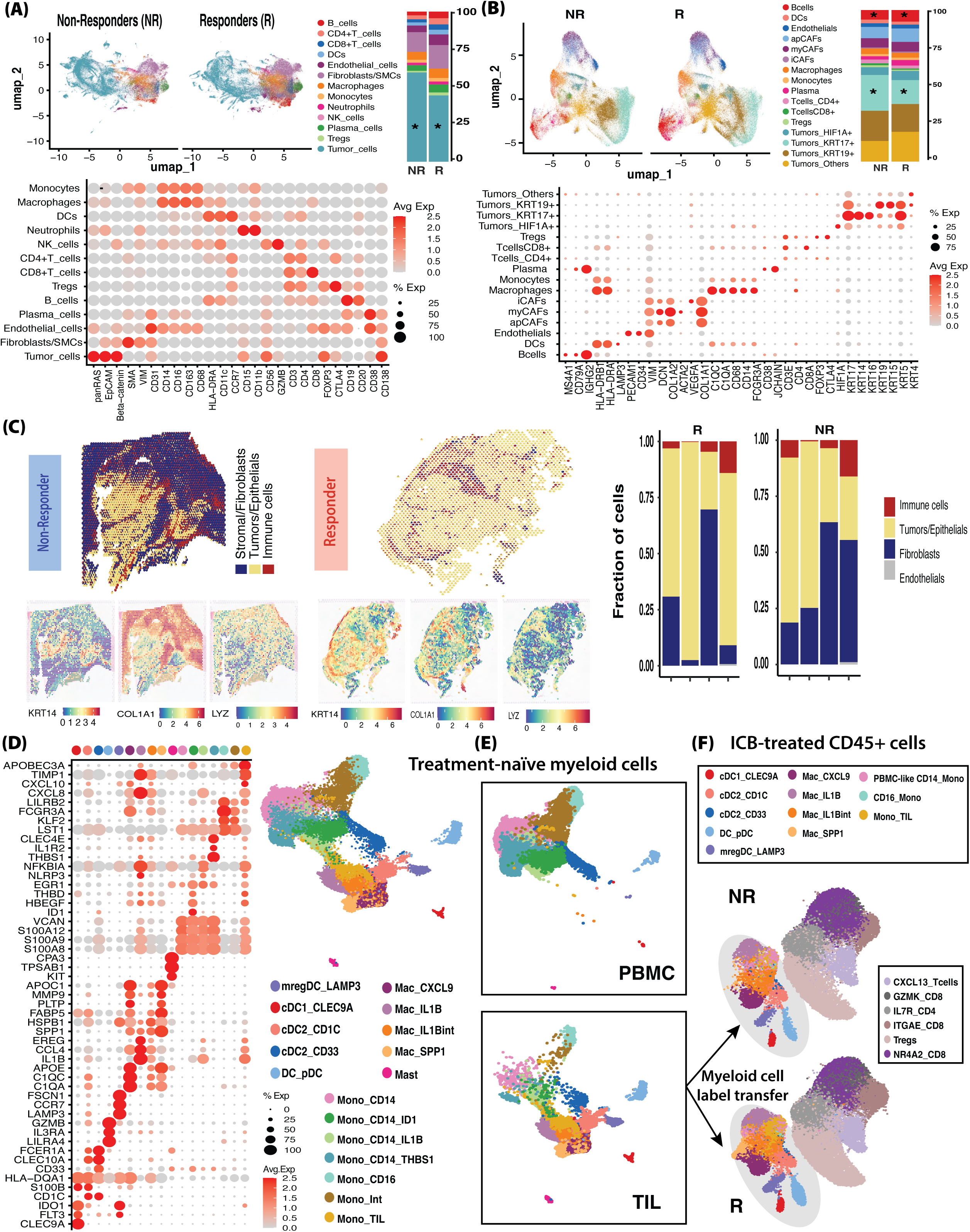
Overview of spatial and single-cell immune landscape in HNSCC treated with Immune Checkpoint Blockade (ICB) a. Uniform Manifold Approximation and Projection (UMAP) visualization, average percentage by cell type (star annotated the cell type abundance difference between Responders (R) and Non-Responders (NR), with p<0.05, Wilcox test, also see **Suppl 1A** for detailed comparison), cell type gene signatures defined from differentially expressed gene (DEG) list, of HNSCC R and NR patients treated with ICB identified from Nanostring CosMx single-cell spatial protein profiling data. b. UMAP visualization, average percentage by cell type (star annotated the cell type abundance difference between R and NR, with p<0.05, Wilcox test, also see **Suppl 1C** for detailed comparison), cell type gene signatures defined from DEG list, of HNSCC R and NR patients treated with ICB identified from Nanostring CosMx single-cell spatial RNA profiling data. c. Major cell type compartment percentage (using computational deconvolution) by spot for representative NR and R samples sequenced with 10X Visium spatial transcriptomics, and representative cell type markers. Distribution of cell type compartment proportion in 8 R and NR samples from 10X Visium. d. Gene signatures from DEGs of different myeloid subsets, UMAP visualization from reanalyzed treatment-naïve HNSCC scRNA-seq. e. UMAP visualization of reanalyzed myeloid subsets split by PBMC and tumor samples. f. Label-transferred defined myeloid cell subsets and established T cell subsets in IBC-treated HNSCC R and NR using computational label transfer.

Among them, we identified a lower abundance of KRT17+ tumor cells in Responders (p=0.02, **Suppl Fig 1C**), as reported previously^16^. In addition, B cell and Plasma frequency trends were higher in Responders (p=0.07 and p = 0.05, **Fig 1B, Suppl Fig 1C**). We also observed increased CD4+ and CD8+ T cells and fewer Tregs in Responders (p>0.05). Notably, the major cell type frequencies correlate highly in two CosMX assays (**Suppl Fig 1D**), and NK cells were not sufficiently detected (<0.2% from protein assay, Suppl Fig 1A, and none from RNA assay).

To complement the small size of TMA cores and the limited number of 1000 RNA transcripts captured in CosMX data, we generated whole transcriptome spatial data using spot-based 10X Visium from FFPE-preserved tumor slides of 8 ICB-treated HNSCC patients (4 Responders and 4 Non-Responders) (**Fig 1C**). We observed less than 20% of immune CD45+ spots using computational deconvolution of the three major cellular components (Tumor/Epithelial cells, stromal cells/fibroblasts, and immune cells, **Fig 1C**), likely due to the tissue and Visium variability. Deconvolution of cell type frequency per spot enabled us to better characterize the distribution of immune cells, epithelial/tumor cells, and fibroblasts within the tissue and suggested no significant differences in their proportion between Responders and Non-Responders while remaining generally consistent with cell type frequencies in our single-cell spatial transcriptomics RNA data (**Suppl. Fig 1E**).

To identify immune cell-cell interactions, with a focus on myeloid-T cells, in response to ICB in HNSCCs, we started with scRNA-seq by reanalyzing multiple scRNA-seq datasets encompassing both treatment-naive and ICB-treated HNSCC patients to look at ligand-receptor (LR) interactions at the single cell level. To determine tumor-specific myeloid cell heterogeneity, we interrogated 26,444 myeloid cells from both scRNA-seq of PBMC and tumors in treatment-naive HNSCC patients^5^ (**Fig 1D**). Including matched PBMC and tumors helped identify 7 monocyte subsets, including one subset only in tumors (**Fig 1E**), 4 macrophage subsets, and 5 DC subsets. Our reanalysis improved the previous identification of HNSCC tumor-infiltrating myeloid cells (**Fig 1E**)^5^. Next, we used Seurat label transfer to identify these myeloid cell types in publicly available scRNA-seq of 19 ICB-treated oral squamous cell carcinoma patients (8 Non-Responders, 11 Responders) (**Methods**). We recovered all treatment-naive myeloid cell types in the ICB-treated samples (**Fig 1F**), with trending differences, including dendritic cell subset elevation in Responders (**Suppl. Fig 1F**). Comparing the originally published annotations from the ICB-treated cohort with our inferred annotations using a Gene Ontology-based method reveals a high correlation between corresponding subsets, especially within the dendritic cell and macrophage compartments (**Suppl. Fig 1G**). We kept the original annotations for T cell subset annotation comparing across ICB-treated patient cohorts (**Suppl. Fig 1H**). In summary, we provide a comprehensive and diverse single-cell and spatial omics data collection of the HNSCC TME from patients treated with ICB, incorporated with scRNA-seq analysis from previously published ICB-treated tumors in an integrated analysis framework to identify potential immune cell interactions in the form of immune neighborhood hubs or LR expression as markers of ICB responses in HNSCC.

### Single-cell spatial omics analysis identifies stroma-immune mixed cellular neighborhoods in Responder patients

Since cellular neighborhoods, defined by the spatial colocalization of different cell types, are associated with ICB response, we used a window-based neighborhood identification method for single-cell spatial data^14^ to identify cellular neighborhoods in the HNSCC TME from CosMX data. From protein data, we identified 7 neighborhoods with distinct cell type compositions, derived by merging the original 20 neighborhoods identified based on the similarity of cell type enrichment within the neighborhood (**Fig 2A, Suppl Fig 2A**), including a tumor epithelial-rich region (Tumor_High neighborhood) significantly more abundant in Non-Responders (p<0.01, **Fig 2B**). Overall, neighborhoods with immune cells are more abundant in Responders (**Fig 2B**), except for the one enriched for CD11b+CD15+ neutrophils (Neutrophil_High neighborhood), which is lower in Responders. Visual inspection with representative immune markers (CD19, CD3E, HLA-DRA-CCR7, GZMB for B, T, and DCs) from CosMX data showed agreement with the corresponding cell type annotations from the RNA profiles in an immune-rich Responder and an immune-desert Non-Responder (**Fig 2B**). With a 1000-gene RNA assay, we identified nine neighborhoods with distinct cell-type compositions, derived by merging the original 20 neighborhoods identified based on the similarity of cell-type enrichment within the neighborhood (**Fig 2C, Suppl Fig 2B**). Assessing neighborhood frequency per FOV by ICB response identified a neighborhood enriched for immune cells, primarily CD8+ T cells, monocytes and DCs (ap_iCAF_Immune_Mixed), including VEGFA-expressing inflammatory-like cancer-associated fibroblasts (iCAFs) similar to those previously reported^17^, and to lesser extent with antigen-presenting CAFs (apCAFs) more abundant in Responders, and a neighborhood enriched for myeloid cell types and KRT17+ tumor cells (Myeloid_K17+_Tumor_Mixed) in Non-Responders (**Fig 2D**). Comparing ICB Response-specific enrichment of each cell type within these neighborhoods suggests concordance in the cell type frequencies (**Suppl Fig 2C-D**). This neighborhood also included co-localization of conventional antigen presenting T cell activation, like DCs, and the newly defined apCAF^18^ in the absence of tumor cells expressed pro-tumoral and ICB-non-responsive KRT17. Looking into this neighborhood, we found higher CD4+ T and Treg cell frequencies in Responders (**Supp.** Fig 2C). Furthermore, differential expression analysis identified higher CXCL10 chemokine expression in DCs from Responders but higher CXCL9 in monocytes from Non-Responders (**Fig 2E**), while its receptor CXCR3 was found higher in CD8+ T and Treg cells within the same neighborhood (**Fig 2F**). Recent work showed that CXCL9/10-engineered DCs could enhance ICB response in lung cancer by activating T cell immunity^19^; however, we do not observe increased CXCL9 expression in Responder DCs in this neighborhood (**Fig 2E**). Assessing the average distance between myeloid senders, T cell receivers, and tumor cells by Response identified monocytes and Tregs closer to tumor cells in Non-Responders in the Tumor_Immune_Tcell_High neighborhood. We also observe a variable proximity of CXCL10+ DCs and CXCR3+ CD4+ T cells to tumor cells, with half of FOVs being closer, in Responders compared to that in Non-Responders in the apCAF_iCAF_immune_mixed neighborhood (**Suppl. Fig 2E**). Together, single-cell spatial omics data revealed differences in cellular neighborhoods not detected through cell type frequency comparisons. These include the stromal-immune cell neighborhoods containing antigen-presenting cells (monocytes, dendritic cells, and apCAF) associated with CD8+ T cells. These neighborhoods are enriched in Responders showing higher co-expression of the CXCL10-CXCR3 ligand-receptor pair in DC-T cell interactions but lower expression of CXCL9-CXCR3 in monocyte-T cell interactions.

**Figure 2.**
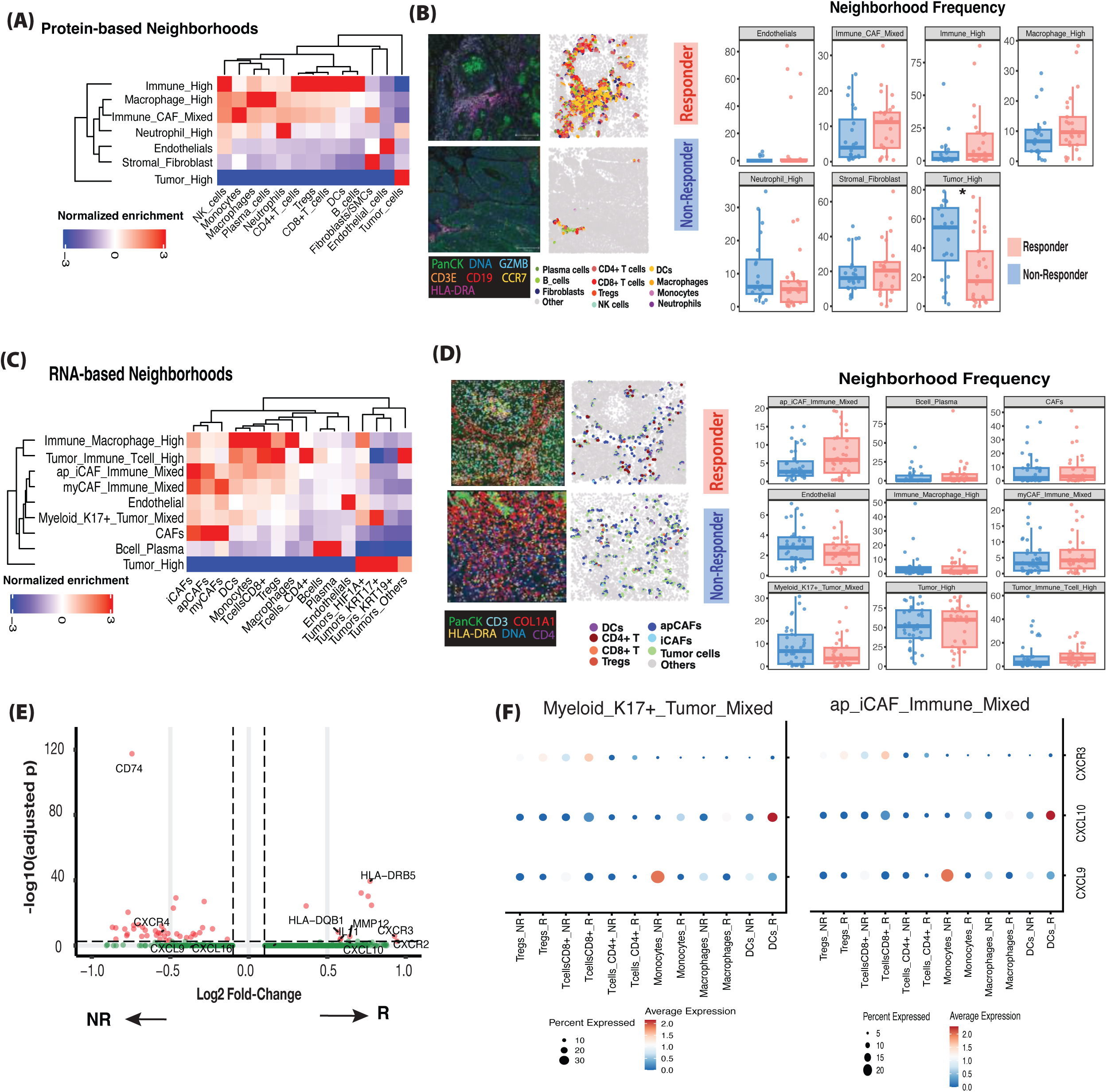
Single-cell imaging-based spatial omics suggests enrichment of a stromal-immune mixed cellular neighborhood in Responders. a. Normalized enrichment score of cell type proportion in cellular neighborhood from protein data identified using a window-based neighborhood identification method^14^. b. Representative fields of view (FOVs) from immune-CAF mixed neighborhood inferred from protein data: the left plot shows representative immune and stromal marker fluorescence and the right plot includes fibroblast and immune cell types within the neighborhood (one R FOV representative above, one NR FOV representative below). Cellular neighborhood frequency comparison between R and NR in protein FOVs. c. Normalized enrichment score of cell type proportion within each cellular neighborhood from RNA data identified using a window-based neighborhood identification method^14^. d. Representative FOVs from immune-CAF neighborhood inferred from RNA data: the left plot shows representative immune and stromal marker fluorescence, and the right plot includes CAF and immune cell types within the neighborhood (one R FOV representative above, one NR FOV representative below). Cellular neighborhood frequency comparison between R and NR in protein FOVs. e. DEGs of DCs in CD45+ neighborhoods between R and NR. f. DotPlot of CXCL9/10 ligand and their receptor (CXCR3) in myeloid and T cell subsets between R and NR in Myeloid_K17+_Tumor_Mixed and ap_iCAF_Immune_Mixed neighborhoods, showing opposite frequency differences.

### ScRNA-seq analysis of myeloid to T cell ligand-receptor interactions in HNSCC response to ICB therapy

To identify myeloid-T cell interactions in HNSCC ICB response, we inferred LR interactions based on the LR CellChat database, with associated biological pathway categorization defined previously^20^ (**Methods**). To compare the difference in Responders vs Non-Responders and before and after treatment, we used the number of LR interactions inferred in each patient cohort (**Fig 3A**) from the scRNA-seq dataset^8^ with our label-transferred myeloid subset annotations from the treatment-naive samples^5^ while keeping and merging original T cell annotations based on GO-based cross-dataset comparisons (**Methods** and **Suppl. Fig 3B**). Various interacting pairs in both Responders and Non-Responders were detected in pre-treatment samples. The primary pre-treatment sender cell types in Responders include IL1B+ macrophages and CD33+ cDC2s, while Non-Responders have various interacting pairs (**Fig 3B**). However, post-treatment interactions include IL1B+ macrophages in Responders and a large increase in significant interactions from PBMC-like CD14+ monocytes and tumor-infiltrating monocytes in Non-Responders to various receiving cell types. Comparing Non-Responders and Responders, fewer interactions were identified in the pre-treatment samples (**Fig 3B**). In Non-Responder patients, we found more interactions in the after-treatment samples, mostly from interactions to all T cell subsets with CD14+ classical monocytes with similar gene signatures to those defined in PBMCs as senders (**Fig 3B**). Notably, of all pre-ICB Non-Responder-specific CD14+ monocyte interactions, 76% remained Non-Responder specific post-ICB, 10% became non-significant, and 13.3% shared. Conversely, of all Responder-specific CD14+ monocyte interactions pre-treatment, 52.3% became Non-Responder specific, 38.6% became non-significant, and 9% became shared (**Fig 3C-D**). For example, Non-Responder interactions conserved in pre- and post-treatment and included chemokine (CXCL16-CXCR6), galectin (LGALS9-CD44, LGALS9-PTPRC, LGALS9-P4HB, LGALS9-HAVCR2), thrombospondin (THBS1-CD47), and costimulatory/coinhibitory (CD86-CTLA4, CD86-CD28) interactions. Lastly, at the per subset level, the one subset where all Non-Responder specific pre-treatment interactions maintain significance and persist as Non-Responder specific post-treatment are the ITGAE+ CD8+ T cells. Narrowing in on enriched sender-receiver pairs at the pathway level showed mregDCs as the top sending cell type post-treatment in both groups in the chemokine pathway, showing little preference per group, which was similarly seen in the whole database analysis (**Suppl Fig 3A).** Within the complement pathway, Responders have an increase in outgoing signaling post-ICB from CXCL9+ Macrophages, whereas most outgoing signaling from this population pre-ICB was found in Non-Responders, but these interactions do not have a big difference as in other pathway categories (**Suppl Fig 3B).** Lastly, outgoing cDC1 signaling within the Costimulatory/Coinhibitory pathway was enriched in Responders, a trend reflected in the whole database analysis post-ICB, but not pre-ICB through an increased number of receiving cell types interacting with cDC1s post-ICB in the Responders (**Suppl Fig 3C).** These data suggest that interactions enriched in ICB treatment, such as CXCL16-CXCR6, THBS1-CD47, and CD86-CTLA4, originating from CD14+ monocytes with a transcriptional signature similar to that of PBMC CD14+ monocytes, are prevalent in post-treatment Non-Responders.

**Figure 3.**
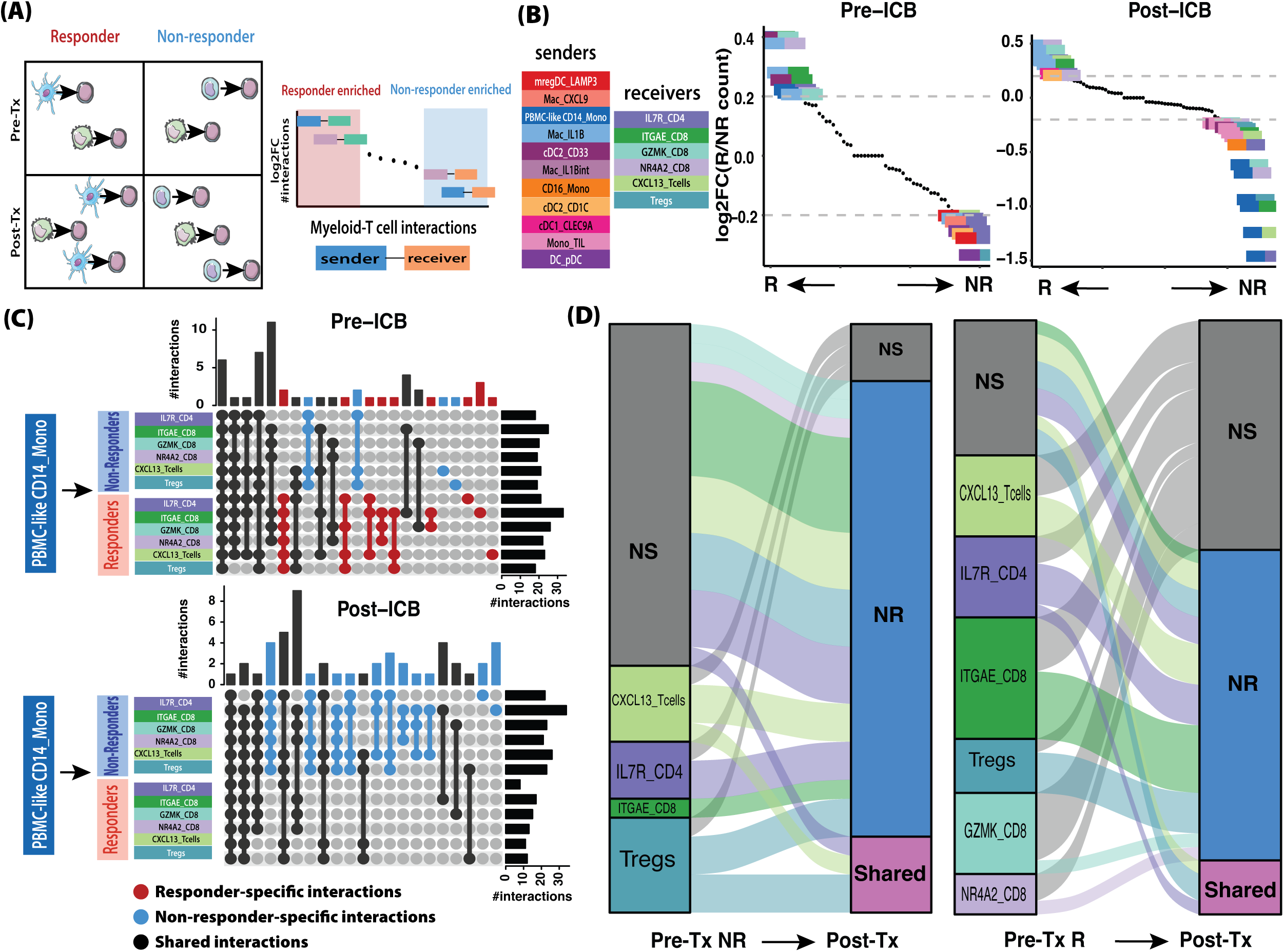
CD14+ monocytes sharing a transcriptional signature with PBMC monocytes have increased outgoing signaling in Non-Responders. a. Schematic of *in silico* cell-cell interaction (CCI) analysis. Comparison analyses were plotted with interaction rank between groups (with R-enriched interactions at the lower end) on the x-axis and log2 fold change of CellChat interaction count on the y-axis. Enriched interactions are visualized with the left box representing senders, and the right box representing receiver cell types. b. Rank plot of CCIs enriched by Response for pre- and post-ICB. c. UpSet plots representing the degree of overlap of LR interactions to receiving T cell types sent from PBMC-like CD14 monocytes both pre- and post-ICB. d. Alluvial plots representing sent signal to T cell subsets from PBMC-like CD14 monocytes, pre-ICB, and the specificity and receiving cell type of these same interactions post-ICB. The left strata and links represent pre-ICB receiving cell type, and the right strata represent post-ICB specificity by Response. Grey stratum represents interactions either gained or lost following ICB. Left: post-ICB specificity of NR-specific interactions from pre-ICB. Right: post-ICB specificity of R-specific interactions from pre-ICB.

### Integrated single-cell and spatial omics analysis refined context-specific immune cell LR interactions in HNSCC ICB responses

Because we found heterogeneity of ligand expression in different myeloid cell subsets **(Fig 2F**) that are spatially and cell-type dependent, we evaluated LR interactions between myeloid and T cells at the levels of both cell-type-specific and spatial resolutions using scRNA-Seq and CosMX data. First, we divided the LR interactions into four functional pathways: Chemokine, Complement, Interleukin, and Co-stimulatory/Co-inhibitory (**Fig 4A**). We identified most interactions pre- and post-ICB from the Costimulatory/Coinhibitory pathway, but also with notably higher pre-treatment interactions through the Complement pathway, especially those “sent” from macrophages in Responders and monocytes in Non-Responders (**Suppl Fig 4A-B**). When comparing paired samples from pre- and post-ICB in Responders and Non-Responders by pathway, we observe varied patterns including enriched pre-ICB chemokine signaling from monocytes and post-ICB chemokine signaling from cDC1 DCs in Non-Responders. In contrast, ICB Responders had enriched post-treatment chemokine signaling sent from mregDCs (**Suppl Fig 4C**). pre-ICB complement signaling in Responders was primarily sent from IL1B+ macrophages, whereas CXCL9+ macrophages were the primary complement signaling sender pre-ICB in Non-Responders (Suppl Fig 5D). Costimulatory and Coinhibitory interactions were numerous and varied greatly by ICB response and treatment stage (**Suppl Fig 4E**). Interleukin pathways, specifically IL10-IL10RA/IL10RB, were found in pre-treatment Non-Responders and mainly from macrophages (**Fig 4A**). Chemokine interactions were sparse but generally shared between Responders and Non-Responders. Among chemokine interactions enriched in pre-treatment Responder samples, we found CXCL16-CXCR6 between CD33+ cDC2s and CXCL13+ and tissue-resident memory CD8+ T cells (CD103/ITGAE+ cells), and CXCL9-CXCR3 between cDC1 to the same two T cell subsets and GZMK+ CD8+ T cells (**Fig 4B**). In contrast, the top enriched interactions in pre-treatment Non-Responder samples were less than 1.5-fold in their changes, including CCL4-CCL5 signaling from CD33+ cDC2s to GZMK+ CD8+ T cells (**Fig 4B**). In contrast, the most enriched interactions in post-treatment Responders were CXCL9/10-CXCR3 from LAMP3+ mregDCs to NR4A2+ CD8+ T cells, indicating a cell-type interaction switch after ICB treatment. Like pre-treatment samples, IL16-CD4 interactions between CD16+ non-classical monocytes or CD1C+ cDC2 dendritic cells with CXCL13 T cells, or cDC1 dendritic cell interaction with NR4A2-expressed CD8+ T cells through CXCL16-CXCR6, which were found in different cell type pairs pre-treatment.

**Figure 4.**
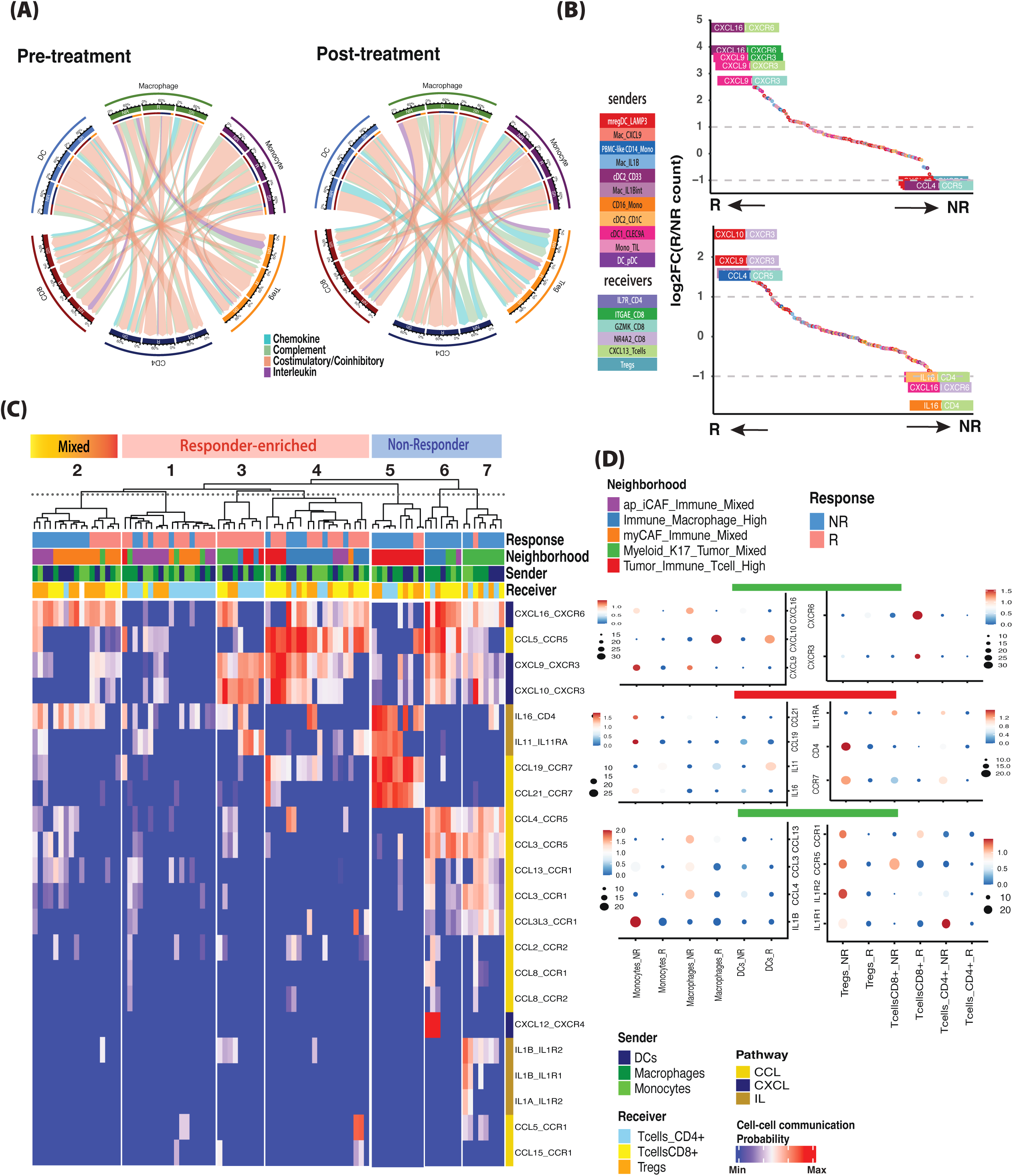
Spatial and cell-type context-specific identification of myeloid-T cell interactions in HNSCC ICB Response. a. Circos plots representing CCI interaction probability from myeloid to T cell compartments by pathway for pre- and post-treatment using previously published pathways^20^. Chord color represents pathway. Interactions from each sender/receiver pair are categorized by group specificity: specific to Responders (R), Non-Responders (NR), or shared (Sh), and converted to percentage of total probability contribution to all interactions within that pair and group specificity. b. Rank plot of CCIs within the chemokine pathway pre- and post-ICB. Interactions on the top left are enriched in R, and interactions at the bottom right are enriched in NR. Individual interactions are represented by a 2-segment box, the left box indicating sender cell type, the right box indicating receiver cell type, ligand is written in the left box, and receptor written in the right box. c. Unsupervised hierarchical clustering identified context-specific myeloid-T LR interaction differences between HNSCC ICB R and NR. (Left) Heatmap represents 7 Ligand-Receptor interaction likelihood probability patterns, merged into 3 major patterns: R-enriched, NR-enriched and Mixed. (Right) LR gene expression in representative neighborhood and cell type specificity identified for R and NR. d. DotPlot for representative LR gene expression in myeloid and T cells within select immune-rich neighborhoods.

To complement and gain better resolution than provided by the scRNA-seq LR interaction analysis, we adapted the CellChat cell-cell interaction (CCI) analysis (**Methods**) for CD45+ myeloid and T cell-rich neighborhoods identified from a window-based cellular neighborhood analysis (**Methods** & **Fig 2C**). Using CCI LR likelihood probabilities, we defined 7 LR interaction patterns using an unsupervised hierarchical clustering method, reflecting context-dependency of spatial neighborhood and cell types of senders and receivers. Furthermore, we merged into three major groups (**Fig 4C**): (i) Responder-specific LRs including CXCL9/10-CXCR3 and CXCL16-CXCR3; (ii) Non-Responder-specific LRs including IL16, IL11 and CCL19/21 signaling in Tumor_Immune_Cell_High neighborhood as well as CCL3/4-CCR5 and IL1B signaling mainly in the Immune_Macrophage_High and Myeloid_K17+_Tumor_Mixed neighborhoods; and (ii) Mixed Responders and Non-Responders with IL16 and CXCL16-CXCR6 signaling in the myCAF_Immune_Mixed neighborhood. Examples were provided in representative contexts, with CXCL10-expressed DC/macrophages interacting with CXCR3-expressed T cells in Responders, CCL19/21-expressed monocytes interacting with CCR7/CD4-expressed Treg and CD4+ T cells in Non-Responders, and CCL3/4/13-expressed macrophages interacting with CCR5-expressed Treg in Non-Responders (**Fig 4D**). Additionally, in the Immune_Macrophage_High neighborhood, we observed CXCL9 most highly expressed in dendritic cells and CXCL16 in macrophages from Non-Responders, and CXCR3 was not highly expressed in any T cell population from these same patients, suggesting that T cell recruitment in this neighborhood may be limited by a lack of CXCR3 in the T cells. In contrast, CXCL10 in DCs and CXCR3 in CD4+ T cells were enriched in ICB Responders (**Suppl Fig 4F**). In the Tumor_Immune_Tcell_High neighborhood, we observed that Responders had enriched CXCL10 in macrophages, CCL5 in DCs, and CXCR3 in CD8+ T cells (**Suppl Fig 4G**). Interestingly, we also observed a group of CCIs, enriched in Responder samples, with CD4+ T cells as receivers mainly from ap_iCAF_Immune_Mixed and MyCAF_Immune_Mixed neighborhoods with few LRs from the three most dominant immune signaling pathways (IL, CCL, CXCL). In summary, our analysis revealed context-dependent myeloid-T cell interactions that correlate with ICB response status in HNSCC ICB treatment. This includes thoroughly examining established interactions (such as CXCL9/10-CXCR3) and several under-appreciated LR pairs that have not been documented in HNSCC and ICB contexts.

### Spatially-constrained immune cell-cell interaction inference identified potential LR interactions enriched in ICB Responders

To further interrogate the spatially-constrained immune-cell interactions enriched in HNSCC ICB response with larger tissues than the TMA cores, we looked at Visium spots from the eight patients’ data (**Fig 1C**) that had at least 5% of immune cells inferred by computational deconvolution (**Methods**). We computed a bidirectional co-expression score to identify which co-expressed LR pairs were enriched by ICB response (**Fig 5A**). Our scoring method identified LR pairs enriched in Responders, including HLA-C-CD8A, CXCL16-CXCR6, CXCL9/10-CXCR3, CXCL13-CXCR3 (**Fig 5B**), CXCL11-ACKR3, DLL-NOTCH1, and TNF-TNFRSF1A (**Suppl Fig 5A-B**). In contrast, Non-Responders enriched interactions included CXCL3-ACKR1, CXCL5-ACKR1, AND CCL13-ACKR1 (**Suppl Fig 5A-C**). The limitation of CD45+ spots and marker expression did not allow performing deconvolution at the resolution of immune subtypes inferred in scRNA-seq; however, we identified CXCL9/10-CXCR3 and CXCL16-CXCR16 for which LR bidirectional scores were enriched in CD45+ spots in tissues (**Fig 5C**). In summary, spot-based Visium data assists in identifying potential candidates for LR pairs, including known markers CXCL9-CXCR3, CXCL16-CXCR6, which were not previously reported, co-expressed in spatially-constrained locations in Responder tissues through a bidirectional co-expression scoring system.

**Figure 5.**
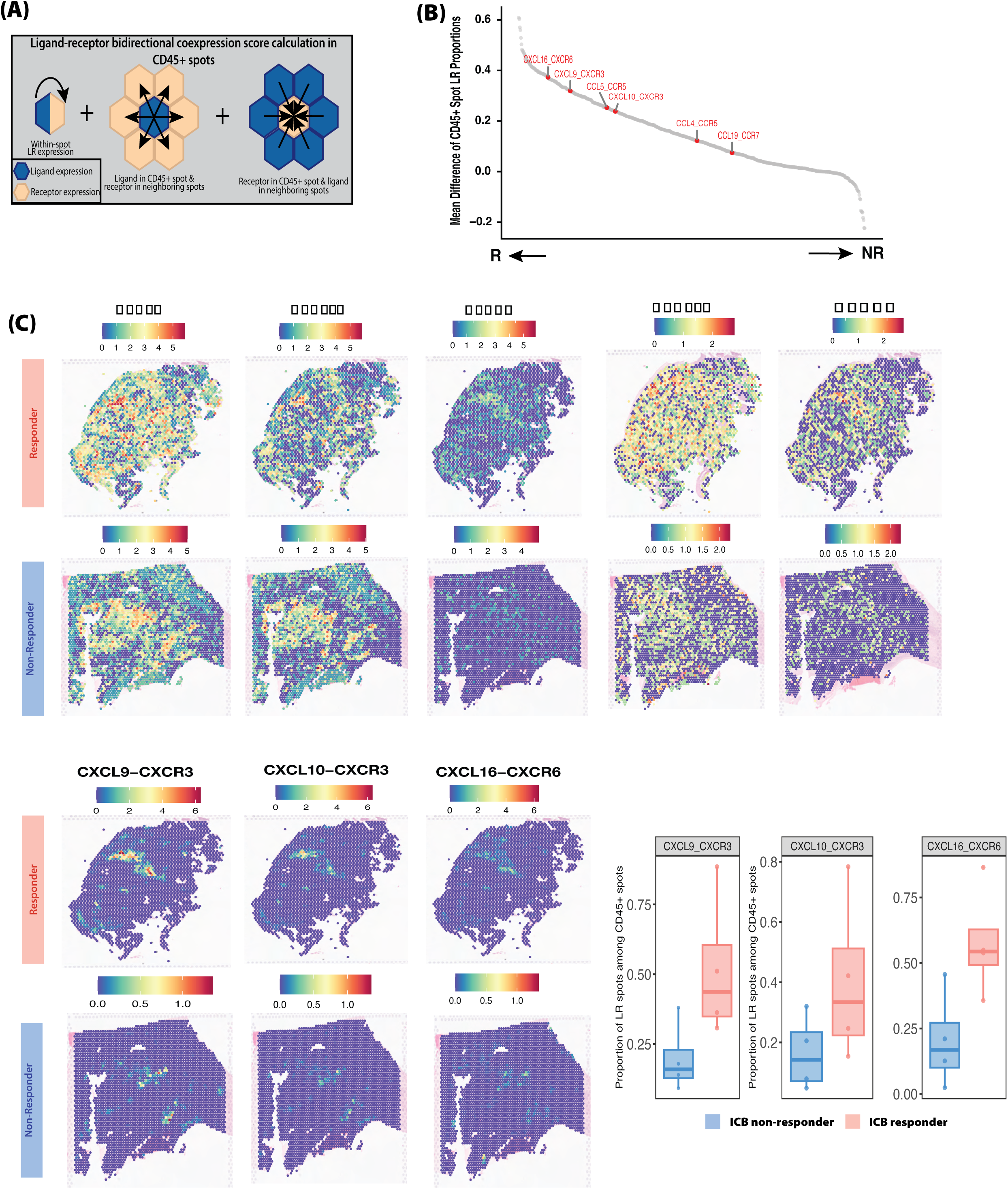
CXCL9/10-CXCR3 and CXCL16-CXCR6 are elevated in immune-rich regions of Responder tumors in Visium tissues. a. Schematic of LR bidirectional co-expression score calculation in CD45+ spots (as defined by >5% immune cells per spot from SCDC deconvolution). b. Rank plot of the difference in R vs. NR mean CD45+ spot proportion for a given LR pair. The left side of the plot highlights interactions enriched in R, and the right side highlights interactions enriched in NR. Select chemokine interactions are highlighted in red. c. Top: Spatial feature plot of CXCL9, CXCL10, CXCR3, CXCL16, and CXCR6 expression per spot for one example R (top) and NR (bottom) tumor. Bottom: Spatial feature plot and boxplots of select bidirectional co-expression LR scores per spot in CD45+ spots for R and NR tumors. The frequency of LR+ CD45+ spots among CD45+ spots was split by response for CXCL9-CXCR3, CXCL10-CXCR3, and CXCL16-CXCR6.

### Evaluating spatial and cell-type context-dependent Ligand-Receptor immune cell interactions using publicly available and independent ICB-treated HNSCC cohorts

To evaluate LR pairs from the Responder and Non-Responder group (**Fig 4C**), we used a publicly available bulk RNA-Seq dataset of CD68+ and CD45+ cells from Nanostring Digital Spatial Profiling technology for an ICB-treated TMA^21^ and a customized panel gene expression of HNSCC patients treated with ICB from a clinical trial^22,23^. Among the LRs defined from single-cell spatial data, we found that the CXCL10 ligand has the highest expression in CD68+ samples (**Fig 6A-B**). Assessing individual LR pairs, we found CXCL10 and CXCL16 show higher expression in CD68+ from Responders (p=0.013, and p=0.025, Wilcox test, respectively) (**Fig 6B**), but their receptors (CXCR3 and CXCR6) show no change in CD45+ samples. A few noticeable differences from LR pairs identified from CosMX data (**Fig 4C**) were observed in this dataset (**Suppl Fig 6**). Additionally, we confirmed that CXCL9 was a better predictor of ICB response than PDL1(CD274) (p =4e-05, p = 0.57, respectively) in a cohort of 107 patients from clinical trials, providing additional support for CXCL9 as an effective predictor of ICB response in large-scale studies, even compared to known clinical markers (**Fig 6C**). In sum, reanalyzing publicly available RNA-Seq data supports CXCL9/10-CXCR3 and CXCL16-CXCR6 myeloid-T cell context-dependent interactions associated with ICB response in HNSCC (**Fig 6D**).

**Figure 6.**
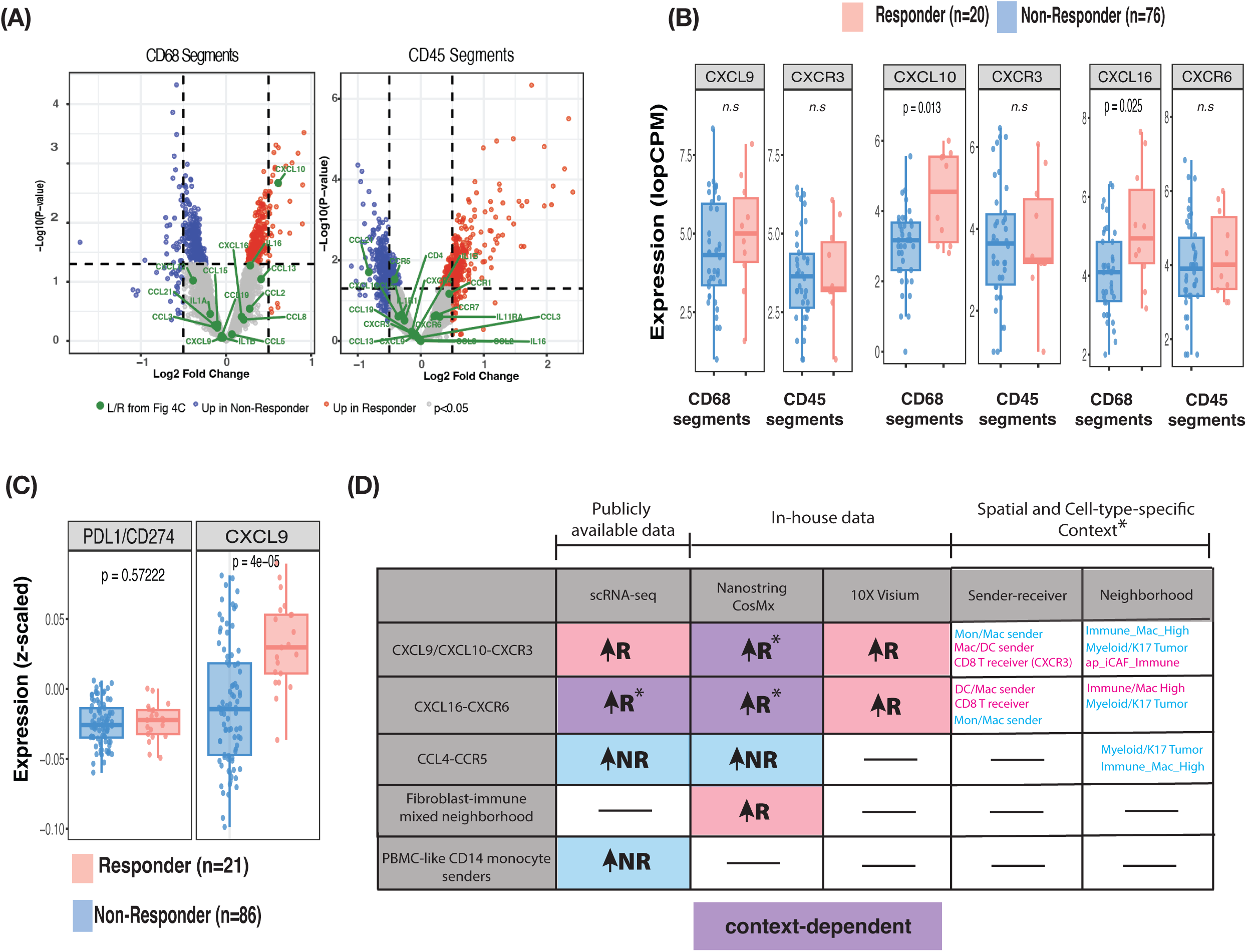
Publicly available independent cohort validation of elevated chemokine interactions in ICB Responder patients. a. Volcano plots show DEGs of CD68+ and CD45+ RNA-Seq from Nanostring GeoMX for NR vs R, highlighting immune LR pairs inferred from **Fig 4C**. b. Comparison of three top LRs between Responders (n=20) vs. Non-Responders (n=76) with Ligands in CD68+ samples and Receptors in CD45+ samples. c. Comparison of CXCL9 and CD274(PD-L1) expression by Response for publicly available clinical trial data from ICB-treated HNSCC patients^22,23^. d. Summary of main differences between myeloid-T cell interactions between HNSCC Responders vs Non-Responders.

## Discussion

HNSCC is the 6^th^ - 7^th^ most common cancer type worldwide. Only 15-20% of HNSCC patients show favorable response to ICB therapy. Improving our understanding of the HNSCC TME may reveal new biomarkers and treatment strategies to augment ICB response and better select those patients most likely to benefit from ICB treatment approaches. In this study, we present a comprehensive, integrated analysis of spatial and single-cell omics of HNSCC TME to evaluate immune cell-cell interactions, focusing on myeloid and T cells in response to ICB. While we did not find much difference in cell type frequencies, despite observing immunological hot tumor phenotypes in Responders^16^, we identified an increase of cellular neighborhoods at the stroma-immune interface in Responders in both protein and RNA single-cell spatial data (**Fig 2**). Interestingly, we identified a decrease in the neutrophil neighborhood in Responders from CosMX protein assay, which was usually underappreciated in spatial RNA assays. Importantly, we defined a spatial and cell-type context-dependent pattern of immune cell-cell Ligand-Receptor interactions in HNSCC response to ICB (**Fig 4C**), including both known and not yet reported interactions in cancer immunotherapy context across different data modalities and profiling resolutions. In addition, we identified an increase of interactions from CD14+ monocytes with T cells in Non-Responders only after ICB treatment from scRNA-Seq (**Fig 6D**).

Our analysis identified multiple candidate interactions in ICB Responders, including widely reported CXCL9/10-CXCR3 between myeloid and CD8+ T cells, an emerging marker of ICB response in HNSCC^7,19,24–28^. In breast cancer mouse models, CXCL9/10+ macrophages were upregulated following dual PD-1/CTLA-4 blockade via a CXCR3-dependent CD8+ T cell infiltration^41^. Our data showed CXCL9/10, mainly expressed in macrophages and DCs (especially LAMP3+ mregDC^29^ subset) and their receptors CXCR3 expressed in CD8+ T cells, associated with ICB response in the absence of CCL4/5-CCR5 interactions between monocytes and CD4+ T cells (including Tregs), highlighting the TME-specificity of immune CCIs. Previous studies have identified both pro- and anti-tumoral roles of CCL4/5-CCR5 in the context of cancer^30^. However, one study in mouse lymphoma and melanoma also revealed pro-tumoral roles of these interactions through the myeloid-T cell axis—identifying CCL4/5 expression in tumor-infiltrating granulocytic and monocytic myeloid-derived suppressor cells (MO-MDSCs), and CCR5 in Tregs. Upon knocking out CCR5, there was a significant decrease in infiltrating Tregs into tumors. In contrast, intratumorally injection of CCL4 or CCL5 increased tumor-infiltrating Tregs, reflecting *in vivo* evidence of our *in silico* observations in the present study^31^. However, we showed a context-dependency of these CCIs due to the complexity and heterogeneous nature of HNSCC tissues in our data. Another interaction, CXCL16-CXCR6, was identified in Responders from scRNA-Seq (between DC subsets and CXCL13+ and tissue-resident CD103+ T cells, **Fig 4B**) and in spatial Visium data (**Fig 5**) but also in Non-Responders in specific cellular neighborhoods from single-cell spatial data (**Fig 4C**). CXCR6 plays an important role in cytotoxic T cells^32^, CD8+ resident memory T cell recruitment in HNSCC^33^. However, CXCR6 was also found to be involved in exhausted T cell phenotypes by a pan-cancer analysis^34^.

Overall, we highlighted the complexity and context-dependent nature of immune cell-cell interactions in human HNSCC, as summarized in **Fig 6D**. This would help further *in vivo* work in mouse models to manipulate and induce these LR interactions for anti-tumoral immunity. Our work also showed the importance of integrated multi-omics analysis in human cancer data, which could generate potential immunotherapeutic biomarkers and targets for HNSCC immunotherapy.

## Declarations

### Ethics approval and consent to participate

The clinical sample collection and analyses were conducted in accordance with the Declaration of Helsinki and received institutional ethics approval from the University of Wisconsin-Madison (UW18144 with IRB#2018–1510, subproject number 2022-009), under a waiver of informed consent.

## Availability of data and materials

All in-house data and source code will be available upon manuscript acceptance.

## Authors’ contributions

HQD, MBF, PFL, and AEGO conceptualized the project. MBF, RH, and TL identified patient samples and generated tumor microarrays to collect Nanostring CosMX and 10X Visium data. AEGO, PK, and HQD analyzed the data. AEGO and HQD wrote and edited the manuscript, with contributions from all co-authors. HQD, MBF, PFL, and PH acquired funding for the project.

## Acknowledgments

The authors would like to thank the patients and their families for contributing precious samples to this project. We acknowledge the Translational Research in Pathology (TRIPath) translational research Biocore at the University of Wisconsin-Madison for tumor microarray assembly and staining for spatial transcriptomics data collection and to the Gene Expression Center (GEC) at the University of Wisconsin-Madison for spatial transcriptomics library preparation and sequencing.

## List of abbreviations

HNSCC: Head and neck squamous cell carcinoma
ICB: immune checkpoint blockade therapy
scRNA-seq: single cell RNA-sequencing
TME: tumor microenvironment
CAFs: cancer associated fibroblasts
DCs: dendritic cells
TMA: tumor microarray
Tregs: Regulatory T cells
CCI: cell-cell interaction
LR: ligand-receptor
R: ICB responders
NR: ICB non-responders
DEGs: differentially expressed genes
UMAP: Uniform Manifold Approximation and Projection
FOVs: fields of view

