## Supplemental Text for "Integrated single-cell and spatial analysis identifies context-dependent myeloid-T cell interactions in head and neck cancer immune checkpoint blockade response"

**Affiliations**

### Materials and Methods

#### Reanalysis of myeloid cells from publicly available treatment-naive and ICB-treated samples

scRNA-seq data from publicly available data was downloaded from NCBI GEO #GSE139324[1] and #GSE200996[2]. Count matrices were used to perform quality control to remove any cells expressing <400 transcripts, <200 or >6000 features. We used single-cell transform (SCT)[3] to regress out mitochondrial and cell cycle genes and normalize gene expression. Principal component analysis (PCA) was performed using the SCT assay, and these PCA values were used to generate UMAP embeddings. Unique clusters were defined by >5 differentially expressed genes using a Wilcoxon rank-sum test with a Bonferroni-corrected p-value < 0.05 and log2 fold change > 0.1. We used Seurat label transfer[4] with default parameters to identify the extent of treatment-naive defined myeloid cell subsets in ICB-treated HNC.

**Comparison of transcriptional similarities between single-cell RNA-seq subsets**

Comparisons of scRNA-seq subsets both within datasets across multiple grouping variables and between datasets were calculated using default parameters with the scGOclust R package[5,] using the GO Biological Processes pathway. The minimum distance used during the analysis was 0.3, and the spread was 0.5.

***In silico* cell-cell interaction (CCI) analyses of single-cell RNA-seq data**

*In silico* CCI analyses of publicly available scRNA-seq[2] were completed using CellChat[6] using the default CellChat human LR database unless otherwise noted. CellChat was run separately on four groups using default settings: Pre-treatment Responders and Non-Responders, Post-treatment Responders and Non-Responders. For comparison analyses, Responder and Non-Responder CellChat objects from the same treatment stage were merged using *mergeCellChat*. CellChat probability scores were extracted using *subsetCommunication* and used for downstream analyses. We compared inferred interacting cell type pairs from Pre- and Post-treatment between Responders and Non-Responders by calculating the log2 fold change of the CellChat interaction count between Responders and Non-Responders and ranking these interactions based on their enrichment in Responders. CellChat scores for Pre- and Post-treatment samples were extracted at the interaction level, subset and aggregated into five pathways using previously defined gene sets[7]. The interferon pathway was removed due to insufficient inferred interactions in the CellChat score results. The interleukin pathway was removed from rank plot analyses where data was too sparse to report.

**Tissue processing for Nanostring CosMX single-cell spatial transcriptomics and proteomics**

A tissue microarray (TMA) was constructed from HNC patients who received ICB-based therapy (Pembrolizumab with or without chemotherapy) at the University of Wisconsin-Madison (n=68 cores from 24 patients) (**Table 1**). To account for tumor and immune heterogeneity within and adjacent to the tumor (i.e., spatial heterogeneity), the TMA was constructed to contain multiple (2-4) tumor sample cores from each tumor, including samples from the central tumor and the leading edge/ TME. A total of 84 x 2 mm cores were included in the TMA. Slides from the TMA blocks were cut by the Translational Research in Pathology (TRIPath) core at the University of Wisconsin-Madison. ICB response was determined using RECIST v1.1 criteria[8].

***Table 1: Patient and tumor characteristics of patients included in the immune check-point blockade-treated head and neck squamous cell carcinoma tissue microarray***

| **Characteristic** | **All patients N = 24** | |
|---|---|---|
| **Primary tumor location** | N | % |
| Oral cavity | 13 | 54.2 |
| Oropharynx | 7 | 29.2 |
| Larynx | 1 | 4.2 |
| Paranasal sinuses | 2 | 8.3 |
| Orbit | 1 | 4.2 |
| **Matched primary + lymph node** | 2 | 8.3 |
| **K17 positive tumor by IHC** | 13 | 54.2 |
| **Best documented response** | | |
| Disease control | 11 | 45.8 |
| Progressive disease | 13 | 54.2 |
| **HPV positive tumor** | 9 | 37.5 |
| **ICB regimen** | | |
| Single agent pembrolizumab | 19 | 79.2 |
| Combination | 5 | 20.8 |

ICB - immune-check-point blockade, IHC - immunohistochemistry, HPV - human papillomavirus

**Nanostring CosMX data acquisition**

Protein and RNA CosMX TMA samples were generated using CosMx SMI by Nanostring Technologies Inc, Seattle, WA[9].Target RNA readout on the SMI instrument was performed following published protocols[9] with a fluorophore-conjugated antibody cocktail against CD298/B2M, PanCK, CD45, and CD3 proteins and DAPI to acquire morphology images. Nine Z-stack images for 5 channels (4 antibodies and DAPI) were captured after washout of unbound antibodies and addition of Imaging buffer. Raw image processing and feature extraction were performed using in-house SMI data processing pipelines[9] that include registration, feature detection, and localization.

**Nanostring CosMX data analysis**

Z-stack images of immunostaining + DAPI were used to draw cell boundaries on the samples. A cell segmentation pipeline[10,11] was used to accurately assign transcripts to cell locations and subcellular compartments. The transcript profile of individual cells was generated by combining target transcript location and cell segmentation boundaries. Cells with fewer than 20 total transcripts assigned were omitted from the analysis.

Normalization using SCTransform (for RNA) and log transform (for protein) and downstream analysis were performed using the Seurat package[4,12–15]. Harmony[16] was used for integration at the FOV level with *npcs = 50*. Graph-based clustering methods were used at different clustering resolutions (0.2 for protein and 0.5 for RNA) and canonical cell type lineages markers were used to annotate the clustering subsets (as shown in **Fig 1A,C**). Cell type frequencies per TMA FOV were calculated. Cellular neighborhoods, based on co-localization of cell-types, were identified from Nanostring CosMX protein/RNA data using the Neighborhood Coordination method implemented in Python[17] using a window size of 10 and k=10. Twenty clusters were initially identified and visually inspected for similar neighborhoods by cell type proportion. Neighborhoods with similar cell type composition were grouped and named according to the enriched cell types in the given neighborhood before being used for downstream analysis, including frequency comparison and cell-cell interaction identification (**Suppl Fig 2A-B**).

Representative morphology and transcript overlay plotting of Nanostring CosMX FOVs was completed using NAPARI (v0.4.17) and the CosMX Tools plugin (Nanostring Technologies, Inc). We used Nanostring-provided NAPARI inputs for both the transcriptomics and proteomics dataset, which both also included morphology staining of cellular compartments and DNA. Pancytokeratin and DNA morphology staining were overlaid onto the tissue, and transcripts of representative markers were then plotted on top of the morphology staining overlay.

**Cell-cell interaction inference from immune-rich neighborhoods in Nanostring CosMX transcriptomics data**

Cell-cell interaction inference from all aggregated immune-rich neighborhoods in Nanostring CosMX transcriptomics data was inferred using CellChat (v.2.1.2) and the full CellChat LR database. Cells from myeloid and T cell rich neighborhoods from all FOVs for each patient were aggregated by ICB response due to the sparse nature of the individual FOVs, and the Seurat GetAssayData() function was used to retrieve SCT normalized values. For both Responders and Non-Responders, the *ComputeCommunProb* function was run with the following parameters: type = 'truncatedMean', trim = 0.1, distance.use = FALSE. Chord diagrams were generated using the *netVisual_aggregate* CellChat function. Ligand-Receptor interaction likelihood probability was extracted from CellChat for hierarchical clustering analysis and heatmap using *ComplexHeatmap* package.

**Spot-based 10X Visium spatial transcriptomic analysis of the HNC tumors**

FFPE blocks from ICB-treated HNC patients treated at the University of Wisconsin Hospitals and Clinics under IRB #2018-1510, subproject number 2022-009, (total: 8 samples, n=4 Non-Responders, n=4 Responders) were selected and processed by the UW TRIPath core at the University of Wisconsin-Madison. FFPE tissue slices were mounted on the 10X Visium capture area with four samples per Visium slide. Mounted samples were incubated, deparaffinized, stained with hematoxylin and eosin (H&E), imaged, and decrosslinked according to manufacturer recommendations by the TRIPath lab. Library construction and sequencing were performed by the Gene Expression Center (GEC) at the University of Wisconsin-Madison Biotechnology Center.

FASTQ outputs from 10X Visium FFPE samples were mapped and pre-processed using SpaceRanger (v1.3.1, 10X Genomics) using the Human Transcriptome Probe set v1. Filtered feature barcode matrices were loaded into R using the *Load10XSpatial* function from Seurat (v4.3.0.1). Spots with zero counts were filtered from the matrix, then data was normalized using SCT. SCT-normalized data was used for PCA, and PCA embeddings were used to identify neighbors and calculate UMAP embeddings using Seurat framework. Cell type inference was performed on Visium data using SCDC (v0.0.0.900)[18] according to the provided vignette using a single cell atlas of immune and non-immune cells from treatment-naive HNC as the reference[19]. Spots with >5% inferred immune cells from this deconvolution were used for subsequent analyses (referred to as CD45+ spots) where indicated to restrict LR interactions to only immune-rich regions of tissue.

**Calculation of ligand-receptor bidirectional coexpression score in CD45+ spots**

To identify potential LR immune cell-cell interactions in Visium data, we calculated a bidirectional co-expression score (**Fig 5A**) for each LR pair within CD45+ spots. For each LR pair, the score for each spot was calculated as the sum of within-spot LR expression, ligand expression in the origin spot and receptor expression in its neighboring spots, and within-spot receptor expression in the origin spot and ligand expression in its neighboring spots based on the function below, where i is the spot of interest and N(i) is the set of neighboring spots. In this case, we define neighboring spots as those directly adjacent to the spot of interest. We define the sending and receiving populations as immune cells, using the sum of immune cell proportions in each spot for $p_{.,S}$ and $p_{.,R}$. $x_{.,l}$ and $x_{.,r}$ are the normalized expression levels of the ligand and receptor respectively.

$$\frac{p_{i,S}\cdot p_{i,R}\cdot x_{i,l}\cdot x_{i,r} + p_{i,S}\cdot x_{i,l}\sum_{j\epsilon N(i)} p_{j,R}\cdot x_{j,r} + p_{i,R}\cdot x_{i,r}\sum_{j\epsilon N(i)} p_{j,S}\cdot x_{j,l}}{1+|N(i)|}$$

The proportion of spots with at least 5% CD45+ with non-zero scores was then calculated for each LR interaction for each sample. For comparisons between Responders and Non-Responders, these proportions were averaged across samples and compared based on fold-change as well as difference in proportion. For visualization on samples, the calculated scores were overlaid for each individual spot. We ranked all LR pairs based on their enrichment in Responders, then filtered to only pairs where both the ligand and the receptor were found in >5% of at least one broad immune cell type from publicly available HNC scRNA-seq data[19].

**Analysis of publicly available clinical trial gene expression data**

Clinical trial gene expression data from ICB-treated HNC patients[20] was retrieved from the meta-analysis, and the same data was included via email request from the authors[21].

**Analysis of publicly available Nanostring GeoMx data**

Publicly available Nanostring GeoMx digital spatial profiling (DSP) data[22] from CD68, CD45, and PanCK segments of HNC tumors was downloaded from GEO #GSE226134 and analyzed by provided ICB response metric (CLINICAL.BENEFIT). Raw data was normalized by counts per million normalization with log scaling.

### Supplemental Figure Legends

Supplemental Figure 1

A. Cell type frequency by ICB response for CosMx protein data
B. Keratin gene expression in tumor cell subsets from CosMx RNA data
C. Cell type frequency by ICB response for CosMx RNA data
D. Comparison of cell type frequency between CosMx RNA and protein datasets, statistics generated using a linear model
E. SCDC-estimated cell type frequency by ICB response for broad cell type compartments from 10X Visium spatial transcriptomics data
F. Cell type frequency by ICB response for publicly available ICB-treated scRNA-seq data
G. Pearson correlation of gene expression profiles between re-annotated treatment-naïve tumor-only myeloid cell subsets and re-annoated ICB-treated myeloid cell subsets, generated using the scGOclust R package.
H. Pearson correlation of gene expression profiles between original ICB-treated T cell annotations and re-analyzed treatment-naïve T cell annotations, generated using the scGOclust R package.

Supplemental Figure 2

A. Normalized cell-type enrichment score across 20 neighborhoods inferred from a window-based neighborhood identification method (NeighborhoodCoordination) for CosMx protein data

B. Normalized cell-type enrichment score across 20 neighborhood inferred from a window-based neighborhood identification method (NeighborhoodCoordination) for CosMx RNA data
C. Left: Normalized cell-type enrichment score for the apCAF_iCAF_Immune_Mixed neighborhood independently calculated and split by ICB response. Right: Within-neighborhood cell type frequency in the apCAF_iCAF_Immune_Mixed neighborhood by ICB response.
D. Left: Normalized cell-type enrichment score for the myeloid_K17+_Tumor_Mixed neighborhood independently calculated and split by ICB response. Right: Within-neighborhood cell type frequency in the Myeloid_K17+_Tumor_Mixed neighborhood by ICB response.
E. Average distance between myeloid and T cell subsets with tumor cells by neighborhood. Myeloid cells were required to have CXCL10 expression > 0, and T cells to have CXCR3 expression > 0 to be included in the analysis.

Supplemental Figure 3

A. Left: Ranking of sender-receiver pair enrichment based on CellChat interaction count pre- and post-ICB within the chemokine pathway. Right: Outgoing signaling from mregDCs to T cell subsets by ICB response both pre- and post-ICB.
B. Left: Ranking of sender-receiver pair enrichment based on CellChat interaction count pre- and post-ICB within the complement pathway. Right: Outgoing signaling from CXCL9 macrophages to T cell subsets by ICB response both pre- and post-ICB.

C. Left: Ranking of sender-receiver pair enrichment based on CellChat interaction count pre- and post-ICB within the costimulatory/coinhibitory pathway. Right: Outgoing signaling from cDC1 dendritic cells to T cell subsets by ICB response both pre- and post-ICB.

Supplemental Figure 4

A. Ranking of sender-receiver LR interactions ranked by enrichment in Responders for complement pathways interactions.
B. Ranking of sender-receiver LR interactions ranked by enrichment in Responders for costimulatory and coinhibitory pathway interactions.
C. Ranking of sender-receiver LR interactions by paired pre- and post-treatment samples (ranked by Pre-treatment enrichment) for chemokine pathway interactions.
D. Ranking of sender-receiver LR interactions by paired pre- and post-treatment samples (by Pre-treatment enrichment) for complement pathway interactions.
E. Ranking of sender-receiver LR interactions by paired pre- and post-treatment samples (by Pre-treatment enrichment) for costimulatory and coinhibitory pathway interactions.
F. Average and percent expression of select ligands (in myeloid cells) and receptors (in T cells) within the Immune_Macrophage_High neighborhood.
G. Average and percent expression of select ligands (in myeloid cells) and receptors (in T cells) within the Tumor_Immune_Tcell_High neighborhood.

Supplemental Figure 5

A. Rank plot of Responder-Non-Responder mean CD45+ spot proportion derived from bidirectional LR co-expression scoring (**Methods**).
B. Proportion of LR+ spots among CD45+ spots for select Responder-enriched interactions found in **Suppl Fig 5a**.
C. Proportion of LR+ spots among CD45+ spots for select Non-Responder-enriched interactions found in **Suppl Fig 5a**.

Supplemental Figure 6

A. Ligand expression in CD68 segments, and receptor expression in CD45 segments by clinical benefit from publicly available GeoMx data from ICB treated HNC patients. Expression values were normalized using cpm normalization with a log transform, then z-scaled for visualization purposes.
