## Supplementary figures and images for "Integrated single-cell and spatial analysis identifies context-dependent myeloid-T cell interactions in head and neck cancer immune checkpoint blockade response"

### Supplemental Figures

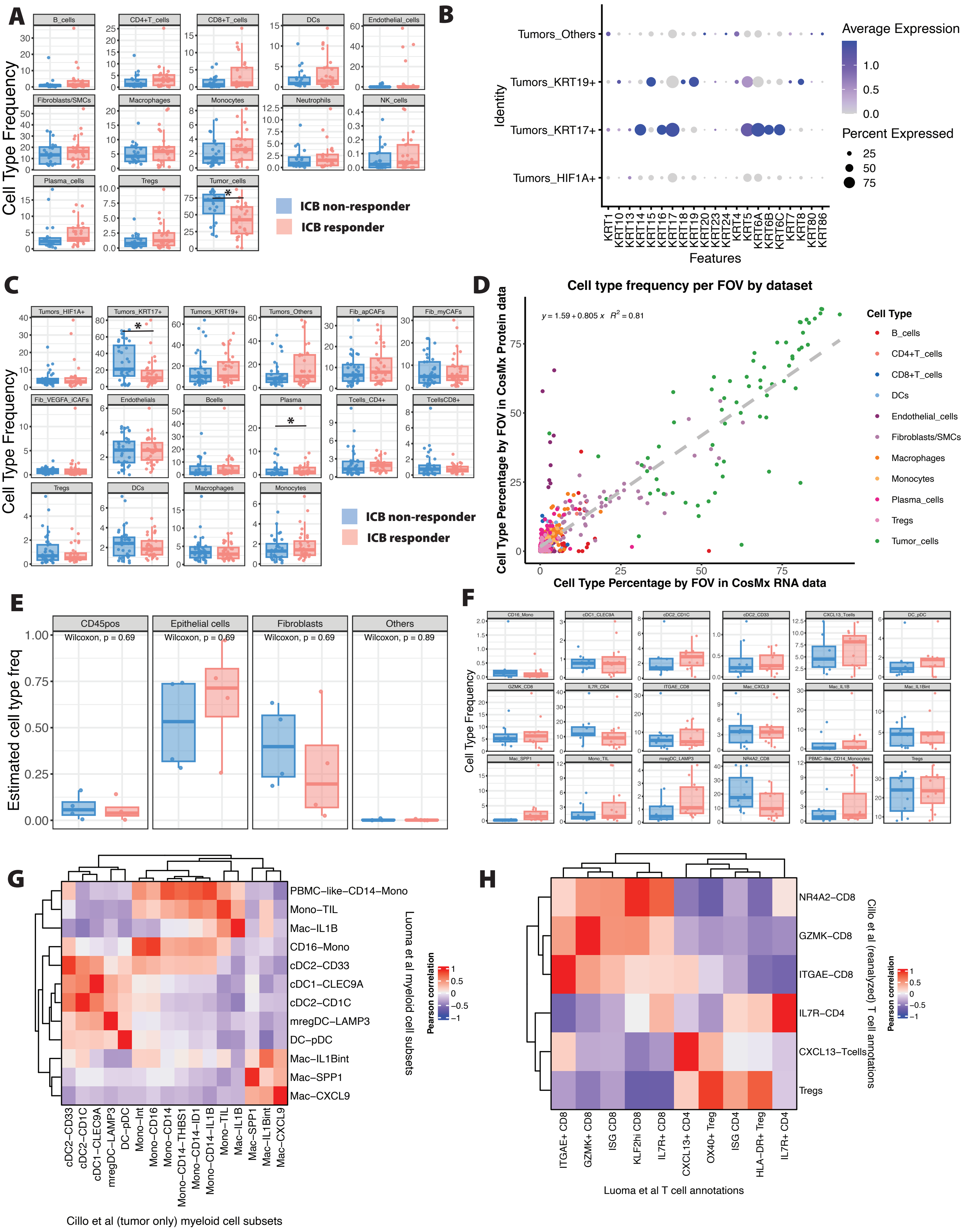

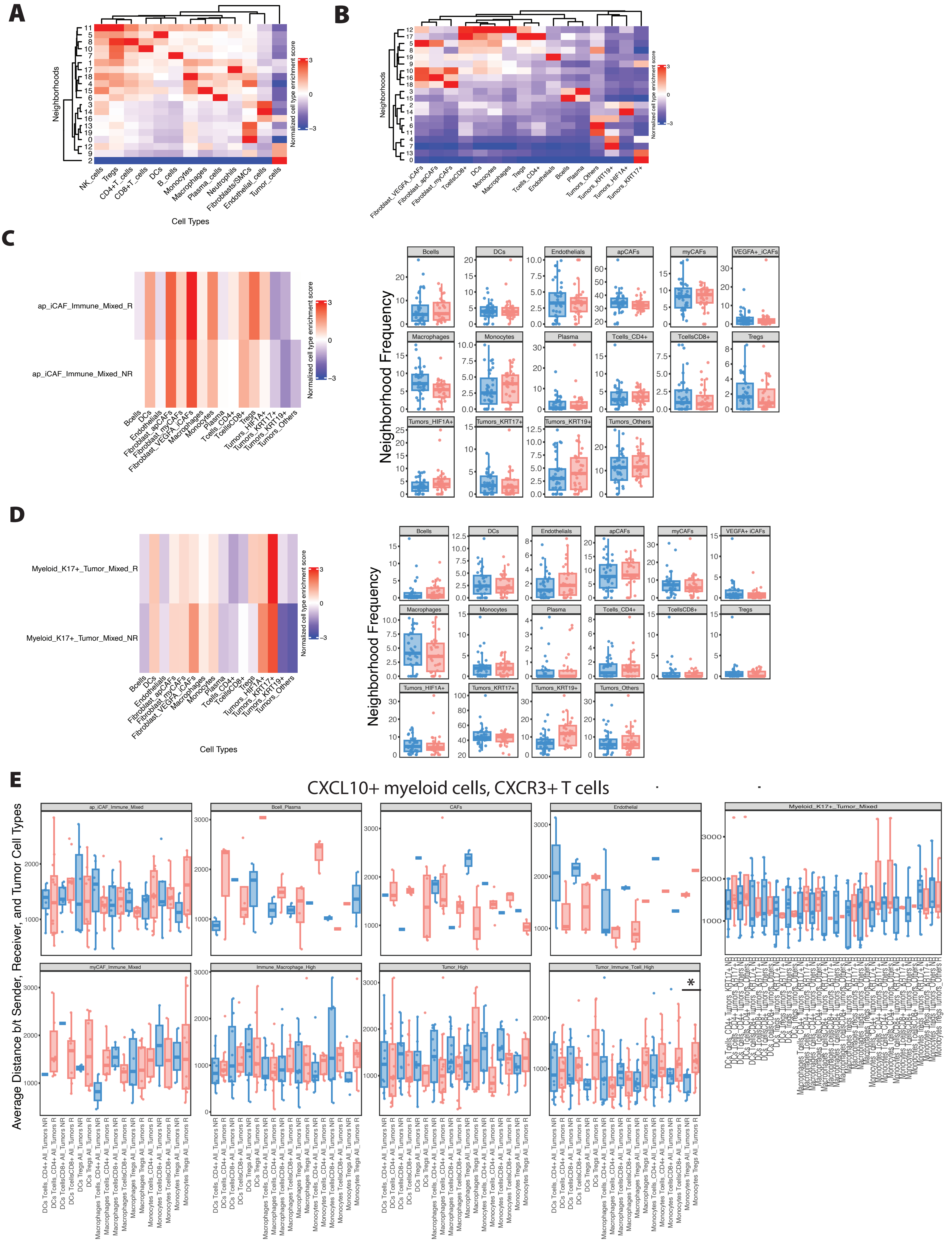

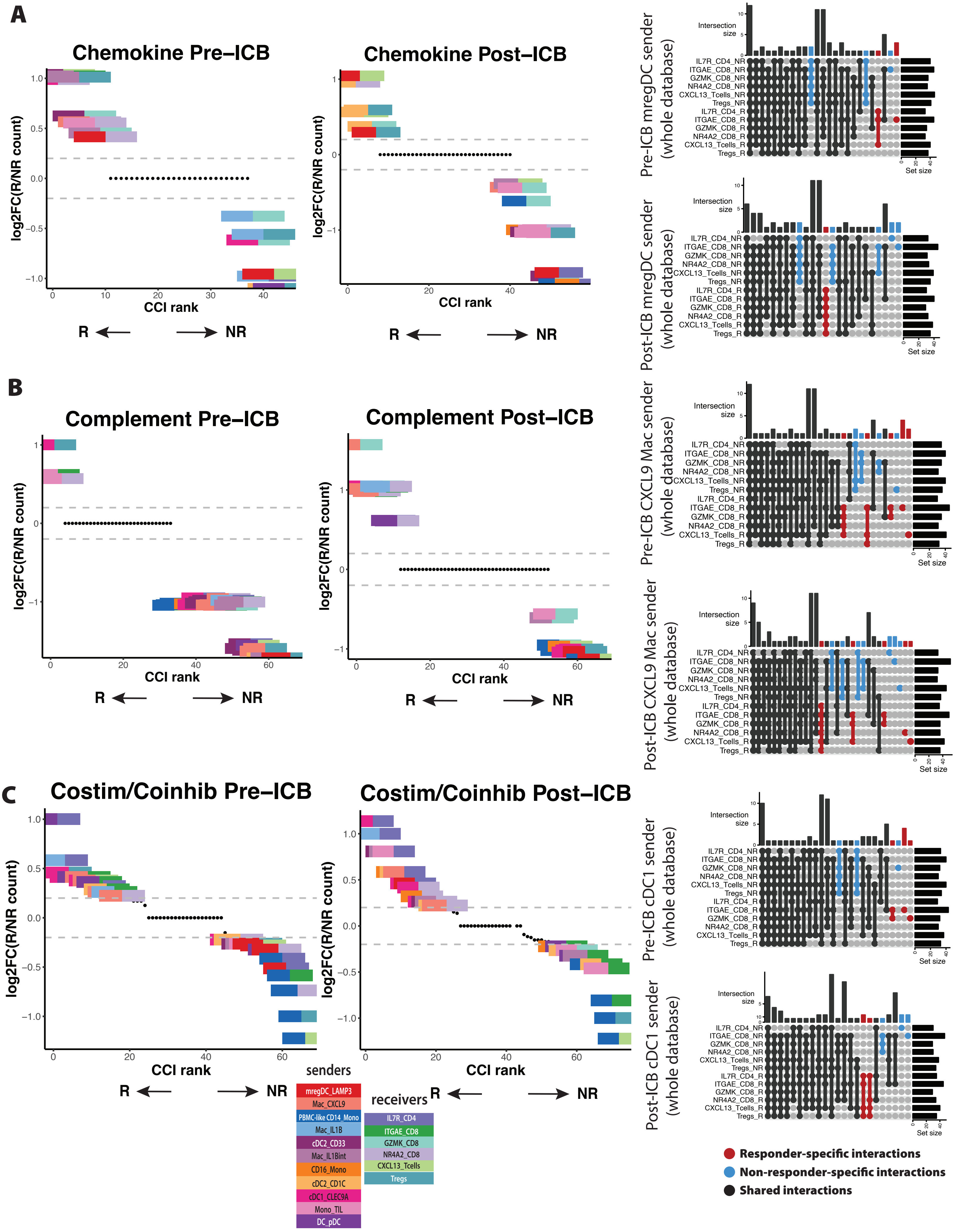

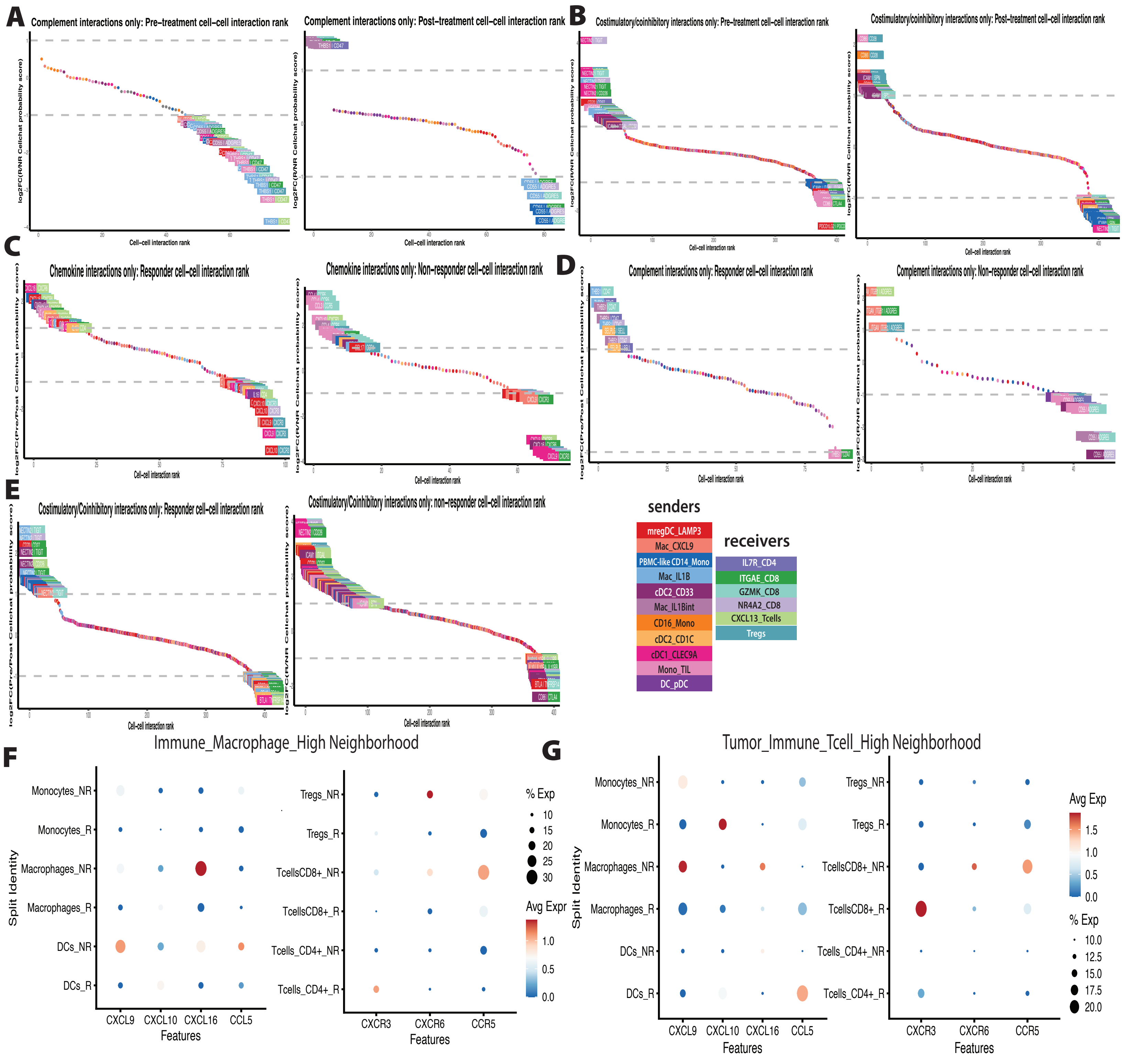

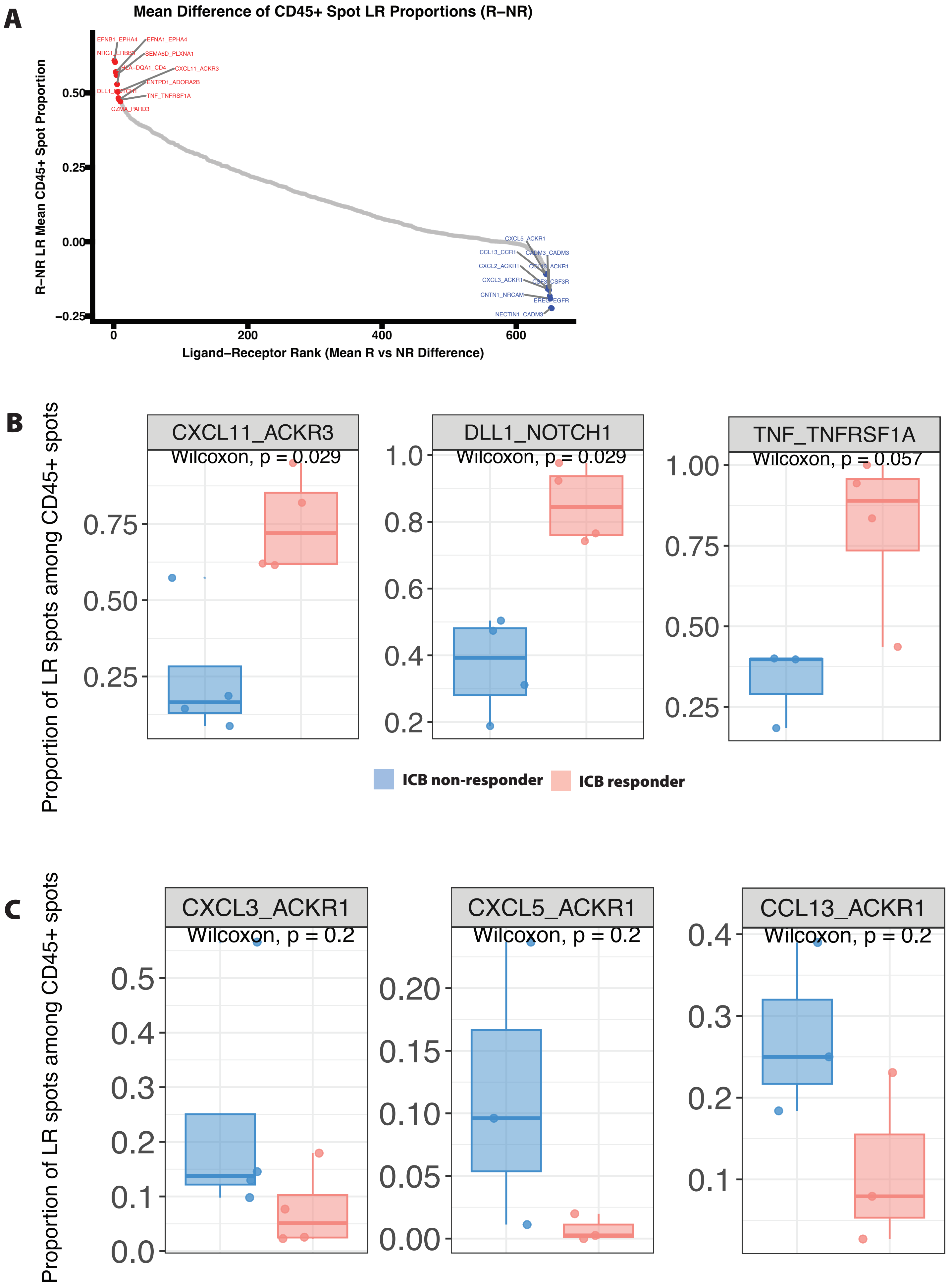

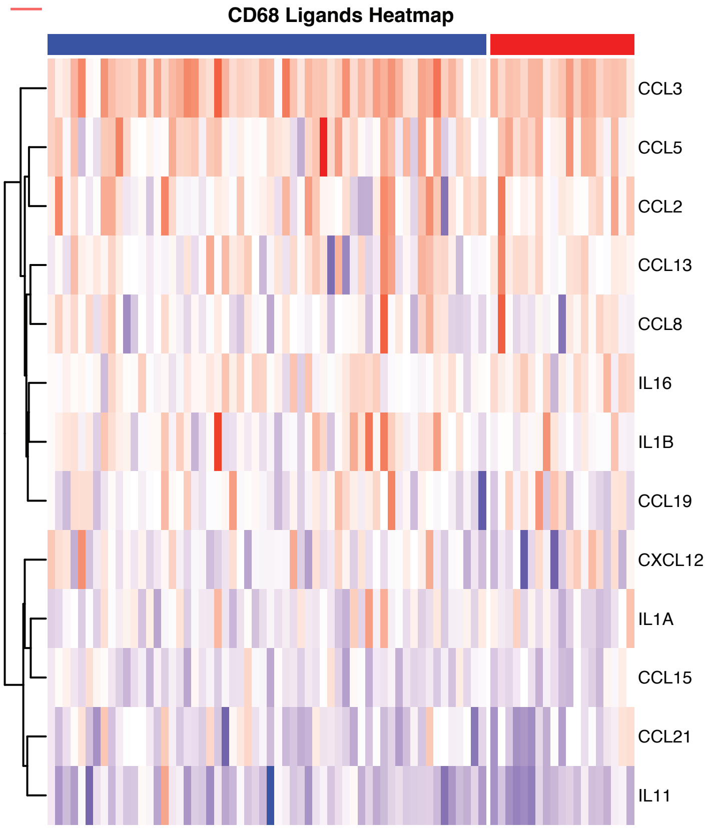

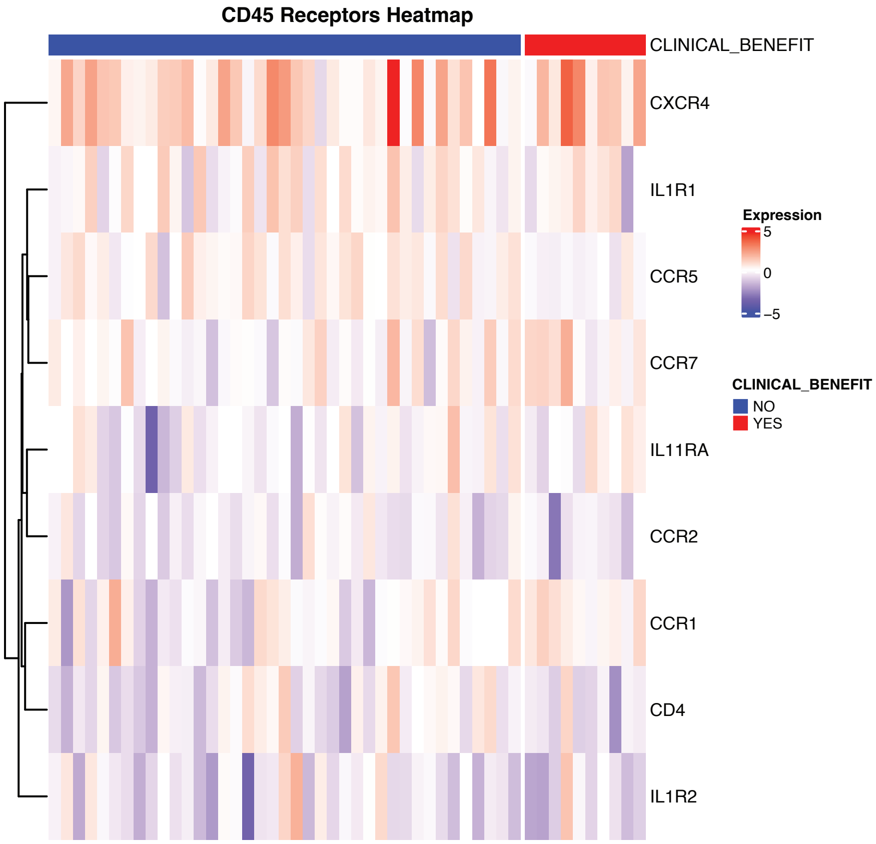
